# Limits of V4 perisaccadic firing rate modulations in explaining perceptual mislocalization

**DOI:** 10.64898/2026.06.16.732392

**Authors:** Geyu Weng, Kelsey Clark, Behrad Noudoost, Neda Nategh

## Abstract

Whether and how various visual sensory areas contribute to the perceived location of visual stimuli remains unknown. To test the role of neurons in extrastriate area V4 in generating alterations in spatial perception during saccadic eye movements (saccades), we examined perisaccadic mislocalization—the perceptual phenomenon in which visual stimuli appearing around the time of a saccade are perceived at a different position than their actual location. We designed and implemented a combined behavioral and electrophysiological framework in non-human primates to directly relate trial-by-trial spatial perception reports during saccades to neuronal firing rates in V4 populations. We measured monkeys’ perception of stimulus location behaviorally and found perisaccadic mislocalization opposite to the saccade direction. We also quantified population responses by computing the center of mass of firing rate activity across probe locations for V4 neurons with receptive fields close to the saccade target. While perisaccadic neuronal responses showed shifts toward the saccade target, these shifts did not systematically vary with the magnitude of perceptual mislocalization across trials. In conclusion, receptive field shifts based on the perisaccadic firing rate of V4 neurons are not sufficient to account for the magnitude of perceptual mislocalization in each trial, suggesting that more complex neural representation of perisaccadic visual information may be critical for linking extrastriate neural activity to saccade-induced perception.

## Introduction

The brain generates a perception of the visual space through processing and interpreting the location of stimuli projected on the retina, but the mechanism by which the brain encodes spatial information is not well understood. To understand the spatial coding of neurons, it is valuable to investigate alterations in neural activity that occur along with changes in visual perception, such as during saccadic eye movements (saccades). Around the time of saccades, visual responses undergo rapid modulation, and our perception of the visual world is altered^1,2^. For instance, our subjective experience of the visual scene remains stable despite the abrupt changes in the retinal image during saccades, a phenomenon known as visual stability^3^.

Another perceptual phenomenon is perisaccadic mislocalization, in which the perceived location of a visual stimulus appearing near the time of a saccade is biased. In total darkness, mislocalization in human subjects starts in the same direction as saccade (forward) and then reverses to the opposite direction (backward), showing a biphasic pattern^4,5^. In monkeys, both forward and backward mislocalization have been observed in darkness^6–8^. Recent studies have reported backward mislocalization for stimuli past the saccade target (ST) in both humans and monkeys^9,10^. In addition to mislocalization in the same or opposite direction to the saccade direction, many studies have reported compression when conducting the experiments in a dimly lit room^11–13^, meaning that stimuli are perceived as closer to the ST (i.e., mislocalization opposite the saccade direction for stimuli past the ST, and in the saccade direction for others). This compression of space is also referred to as convergent mislocalization. The opposite phenomenon is called expansion or divergent mislocalization, in which stimuli are perceived as farther from the ST (i.e., mislocalization in the saccade direction for stimuli past the ST, and opposite the saccade direction for others). The inconsistency in the perceptual reports may be due to differences in paradigm design, such as the ambient lighting, stimulus size or contrast, duration of stimulus presentation, length and direction of the saccade vector, distance between stimulus locations, range of possible stimulus positions, and distance from the stimuli to the ST.

Although numerous studies have characterized the behavioral aspects of perisaccadic mislocalization, its neural mechanisms remain poorly understood. Many studies have investigated the neurophysiology of perisaccadic visual representations in nonhuman primates, and many of them have shown that neurons shift their spatial sensitivity perisaccadically^14–21^. Some neurons in the extrastriate visual areas and prefrontal cortex show a sensitivity shift to the postsaccadic receptive field (RF) even before the saccade, a phenomenon referred to as future field remapping or forward remapping^22,23^. Another type of perisaccadic change in sensitivity, called ST remapping or convergent remapping, is when neural RFs shift towards the ST around the onset of saccade^24–33^. Both future field and ST remapping can be observed in extrastriate visual cortex in the same experiments^34–37^.

Some studies have established computational frameworks based on neuronal recordings to study perisaccadic phenomena^14,36,38–40^. For example, Krekelberg and colleagues used a decoder based on neural data from the middle temporal area (MT), medial superior temporal area (MST), ventral intraparietal area, and lateral intraparietal area in nonhuman primates. Their predictions suggested forward mislocalization near the fixation point (FP) and backward mislocalization near or beyond the ST^14^. Other researchers have also suggested that RF remapping underlies perisaccadic mislocalization^14,41,42^, and some studies have used computational approaches to predict perisaccadic perception of space based on neural responses^42–45^; however, there is a lack of studies linking behavioral reports to neural activity on a trial-by-trial basis. Additionally, how remapping might affect the perceived location varies qualitatively based on whether the same decoding algorithm is applied during the perisaccadic and fixation periods (‘unaware’ of the RF shift) or with knowledge of the perisaccadic response changes (‘aware’)^46^. Comparisons of decoding under these two paradigms have demonstrated that the underlying assumption significantly influences the decoding outcome^46,47^. Future field remapping could lead to no mislocalization when the decoder was ‘aware’ of receptive field shifts; in contrast, the ‘unaware’ decoder produces either backward mislocalization, or biphasic (forward, and then backward) mislocalization under some additional assumptions^47^. For ST remapping, the ‘unaware’ decoder predicted divergent mislocalization, while the ‘aware’ decoder predicted convergent mislocalization^46^. In a previous study, we found spatial bias opposite to the saccade direction around the ST, based on analysis of a statistical model capturing the perisaccadic dynamics of extrastriate spatiotemporal sensitivity^48^. Motivated by the results of this study, we specifically wanted to examine if neuronal modulations around the ST explain behavioral mislocalization on a trial-by-trial basis. Through combined behavioral and physiological experiments, we aim to determine which of the following three hypotheses is best supported by the data: 1) Divergent mislocalization, consistent with ‘unaware’ decoding of V4 activity during ST remapping; 2) Convergent mislocalization, consistent with ‘aware’ decoding of V4 activity during ST remapping; 3) Perisaccadic mislocalization cannot be predicted by the sensitivity shift of V4 neurons alone. We recorded neuronal activity in area V4 as monkeys performed a task to measure the perceived location of stimuli appearing during fixation versus perisaccadically. Behaviorally, we found perisaccadic mislocalization opposite to the saccade direction for stimuli near the ST; however, when trials were divided based on mislocalization magnitudes (‘High’ versus ‘Low’), there was no difference in the firing rate modulation corresponding to the two groups. We then analyzed the spatial profile of the population responses by calculating the center of mass of firing rate responses at nine probe locations; we found that the center of mass of High trials shifts closer to ST during the perisaccadic period, but there is no significant difference between High and Low trials. This lack of difference between High and Low trials was confirmed when restricting the analysis to the probe locations with the strongest behavioral mislocalization. Therefore, we conclude that perceptual mislocalization cannot be accounted for by modulations of V4 perisaccadic firing rates alone, suggesting the involvement of more complex neural coding mechanisms and motivating further investigation into alternative characterizations of saccade-induced response modulations in V4 and other areas.

## Results

A previous study theorized that whether the same decoding algorithm is applied during the perisaccadic and fixation periods (‘unaware’ of the RF shift) or with knowledge of the perisaccadic responses changes (‘aware’) might cause two different types of mislocalization^46^. We adapted the results from this theoretical framework to schematically illustrate the two cases under ST remapping with 25% attentional enhancement and a center-of-mass decoder (Fig. 1a). In the ‘unaware’ case, the decoded population response of neurons will result in divergent mislocalization (Fig. 1a left). In the ‘aware’ case, the decoded population response of neurons will result in convergent mislocalization (Fig. 1a right).

**Figure 1.**
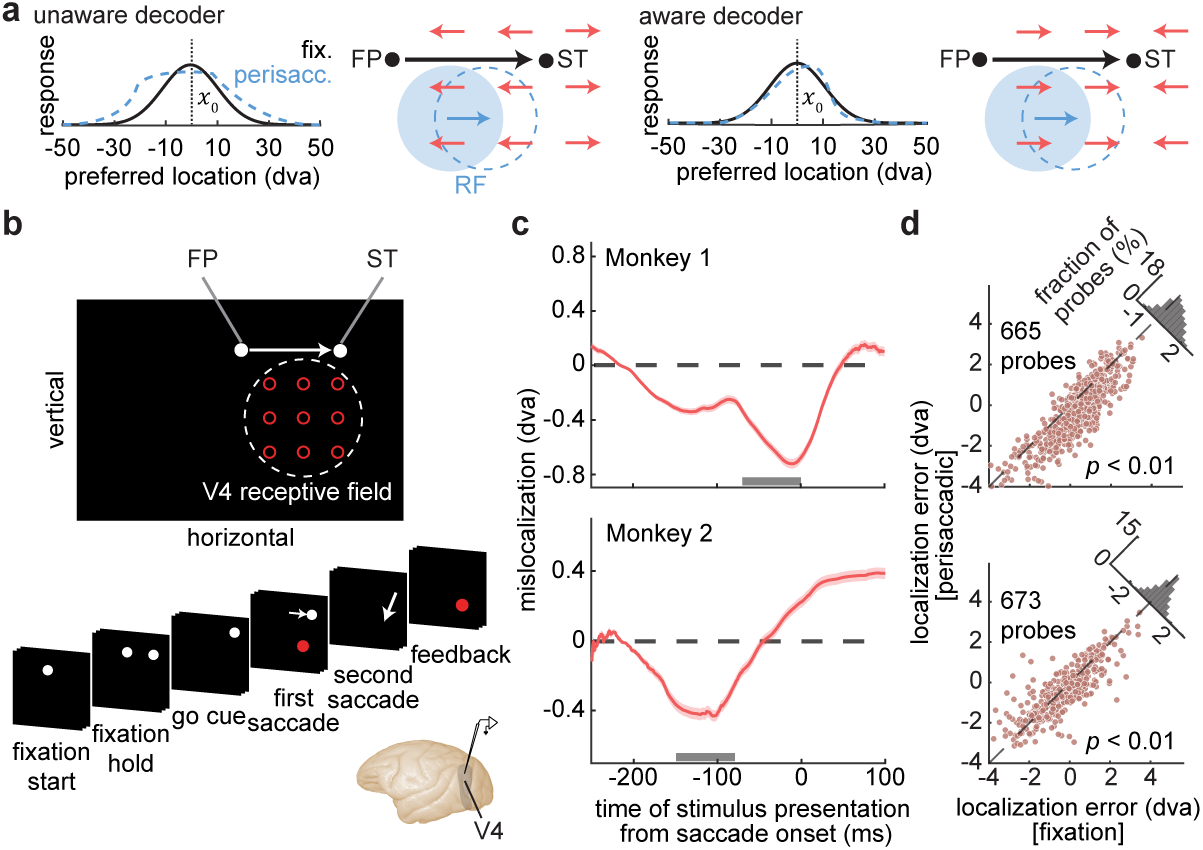
Theoretical hypotheses, experimental paradigm, and behavioral outcomes for perisaccadic mislocalization phenomenon. **a.** Schematics of differing perisaccadic mislocalization resulting from the same neural shift based on the decoder. Perisaccadically, receptive fields (RFs) close to the saccade target (ST) shift convergently toward the ST (10 dva). Assuming the preferred stimulus location (*x*_0_) of the population of neurons is 0 dva, the decoded population response is therefore largest at 0 dva during fixation (black solid line). If the perceptual decoder is ‘unaware’ of the RF shift, the decoded preferred location of population response shifts divergently, away from the ST (blue dashed line; left). If the perceptual decoder is ‘aware’ of the RF shift, the decoded preferred location of population response shifts convergently, toward the ST (blue dashed line; right). As a result, behavioral mislocalization is divergent under ‘unaware’ condition (left) and convergent under ‘aware’ condition (right). The start of the red arrows indicates actual locations of the visual stimuli, and the end of the red arrows indicates the perceived locations when the stimuli are presented perisaccadically. Blue circles indicate an example V4 RF (shaded, fixation; dashed, perisaccadic), which shifts toward the ST perisaccadically (blue arrow). **b.** Perisaccadic Localization Task. In each trial, the monkey makes a first saccade from an FP to a peripheral ST 10 dva away from the FP horizontally (left or right, one ST location per session). At a random time during fixation and saccade execution, a 50-ms visual probe stimulus is presented in one of nine possible locations in a 3×3 grid placed around the V4 neuron’s receptive field. When the ST disappears, the monkey makes a second saccade to the remembered location of the probe stimulus. If the monkey reports within a window of the correct location, the stimulus reappears as feedback. **c.** Mislocalization is measured as the horizontal distance between the reported location of each stimulus when presented perisaccadically compared with when presented during fixation (−250:−180 ms). The plot shows mean mislocalization, and the shaded area represents the standard error of the mean (SEM) for Monkey 1 (top) and Monkey 2 (bottom). The x-axis shows the time of stimulus presentation from saccade onset. The line plot shows the mean mislocalization for trials with stimulus presentation appearing within ±25 ms of the plotted time value; the number of probes included varies for different time points. Positive values indicate mislocalization in the saccade direction, and negative values reflect mislocalization opposite to the direction of the first saccade. The gray bars (Monkey 1: −70:0 ms; Monkey 2: −150:−80 ms) show the perisaccadic time windows used for (d). **d.** Horizontal localization error for probes presented in fixation versus perisaccadic windows. The horizontal localization error is measured as the horizontal distance between the reported location of each stimulus and the actual location of the probe presented (negative values indicate error in the direction opposite the saccade). Plots show horizontal localization error for probes appearing perisaccadically (y-axis) versus during fixation (x-axis) for Monkey 1 (top; p = 3.72e−59) and Monkey 2 (bottom; p = 5.00e−07). p-values were computed by two-sided Wilcoxon signed-rank tests. Histograms in the upper right show the distribution of differences.

We used a combined behavior and neurophysiological paradigm to test whether perisaccadic remapping in V4 neurons explains perisaccadic mislocalization, and to evaluate whether the data support any of the scenarios outlined in Figure 1a. We designed a behavioral paradigm called the Perisaccadic Localization Task to measure perisaccadic mislocalization while recording neuronal activity in area V4 using array or single electrodes (see Methods). Two monkeys performed the task while their eye movements were monitored with a high-resolution optical eye-tracking system. The monkey made a first saccade from the FP to the ST 10 degrees of visual angle (dva) left or right from the FP horizontally. A probe stimulus appeared for 50 ms at one out of nine possible locations at a random time during fixation or around the time of the first saccade (Fig. 1b). After landing on the ST, the monkey then made another saccade to the probe location, and the endpoint of that saccade was considered as the reported location. Because the saccade vectors were horizontal, we defined mislocalization as the horizontal distance between the reported location of each stimulus when presented around the time of the first saccade compared with the average reported location for stimuli presented during fixation—180 ms to 250 ms before the onset of the first saccade (−250:−180 ms). In other words, we calculated the mislocalization for each trial as the reported location of that trial minus the mean reported location across all fixation trials. Positive values indicate mislocalization in the saccade direction, and negative values reflect mislocalization opposite to the direction of the first saccade.

We observed perisaccadic mislocalization opposite to the direction of the saccade. The average time course of mislocalization shows the greatest mislocalization for probes presented shortly before saccade onset (Fig. 1c), and we used the time windows with the greatest mislocalization (Monkey 1: −70:0 ms; Monkey 2: −150:−80 ms; shown by gray bars in Fig. 1c) as the perisaccadic time windows for subsequent analysis, including in Fig. 1d. We also measured the localization error: the distance between the reported location of each stimulus and the actual location of the probe presented. The horizontal localization error for probes presented perisaccadically was significantly more negative—i.e., larger error opposite the saccade direction—compared with those presented during fixation for both monkeys (Fig. 1d; Monkey 1: fixation = 0.23±0.05 dva, perisaccadic = −0.35±0.06 dva, *p* = 3.72e−59, Monkey 2: fixation = −0.35±0.06 dva, perisaccadic = −0.54±0.06 dva, *p* = 5.00e−07; mean ± standard error of the mean (SEM)). Averaging across the localization errors across probes in each session, we observed significant horizontal localization error for perisaccadic probes compared to fixation (fixation = 0.11±0.11 dva, perisaccadic = −0.22±0.10 dva, *p* = 8.21e-17; Supplementary Fig. 1b), but no significant vertical localization error (fixation = 0.30±0.08 dva, perisaccadic = −0.30±0.09 dva, *p* = 0.08; Supplementary Fig. 1c). The fixation and perisaccadic time windows are consistent for all analyses in this study.

To examine the relationship between perceived location and neural responses across trials, we classified trials as High (associated with mislocalization magnitudes in the top 45^th^ percentile) and Low (associated with mislocalization magnitudes in the bottom 45^th^ percentile). We assessed the corresponding neuronal modulations of the two groups using a modulation index, defined as:

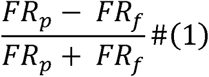

where *FR_p_* represents the firing rate of V4 neurons for probes in the perisaccadic time window, and *FR_f_* represents the firing rate of V4 neurons for probes in the fixation time window across all 9 probe locations. Figure 2a shows the modulation index of High and Low trials over time from stimulus onset. We averaged the modulation index of High and Low trials 60:120 ms from stimulus onset (gray bar in Fig. 2a; chosen based on the time of perisaccadic modulations reported in prior studies^36,48^) and found no difference between the two groups (Fig. 2b; Monkey 1: n = 494, High = 0.04±0.01, Low = 0.04±0.01, *p* = 0.08; Monkey 2: n = 509, High = 0.04±0.01, Low = 0.04±0.01, *p* = 0.94).

**Figure 2.**
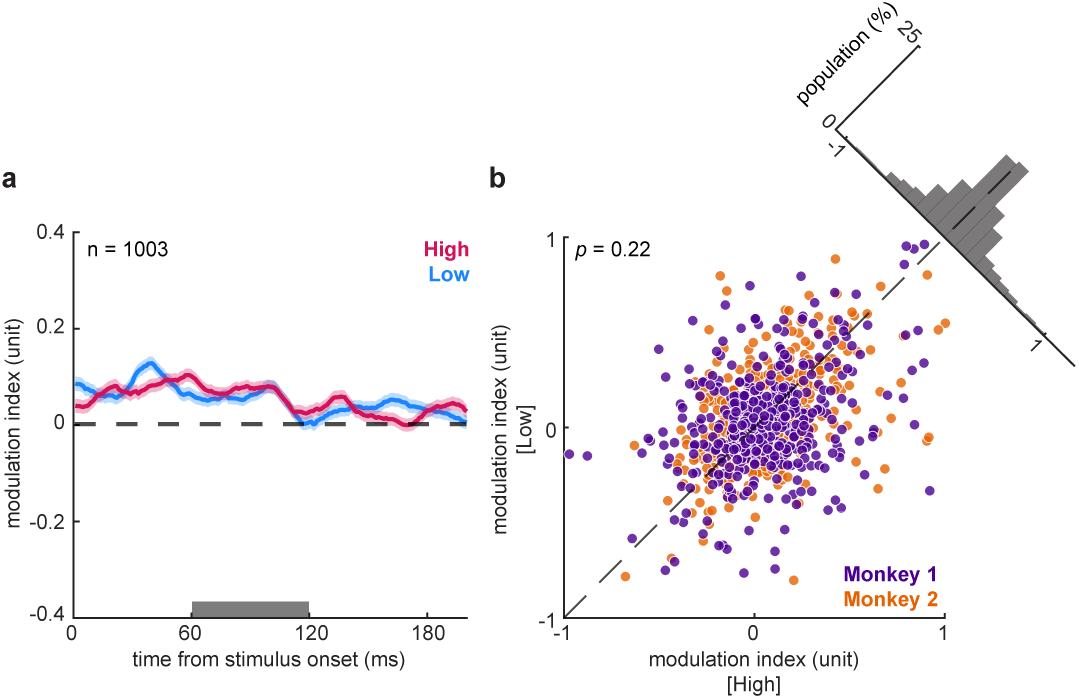
Perisaccadic neural response modulation in trials with higher versus lower mislocalization magnitudes. **a.** The modulation index of High (associated with mislocalization magnitudes in the top 45^th^ percentile; magenta) trials and Low (associated with mislocalization magnitudes in the bottom 45^th^ percentile; blue) trials over time from stimulus onset. Mean ± SEM across neurons. The number of neurons is indicated by ‘n’. The gray bar (60:120 ms) shows the analysis window used for (b). **b.** The mean modulation index within the analysis window for High trials versus Low trials. Plots show the combined population of Monkey 1 (purple: n = 494 neurons, p = 0.08) and Monkey 2 (orange: n = 509 neurons, p = 0.94). p-values were computed by two-sided Wilcoxon signed-rank tests. Histograms in the upper right show the distribution of differences.

Next, we examined whether there is a shift in the spatial sensitivity of the neurons and, if so, whether the shift is related to the magnitude of mislocalization. To examine neurons’ spatial sensitivity shift relative to the ST, we selected neurons with an RF close to the ST (distance between RF center and ST smaller than 15 dva). In the population of 295 neurons from Monkey 1 and 364 neurons from Monkey 2, the center of mass of the neuronal responses shifted toward the ST (Supplementary Fig. 2). To visually illustrate this shift, Figure 3a shows the mean normalized firing rates of High and Low trials of two sample ensembles of neurons (n=11 and n=20 neurons) at nine probe locations for probes presented during the fixation and perisaccadic windows. An ensemble was composed of neurons with similar RF centers that were recorded in sessions with the same probe array position. Both ensembles generally show shifts of the center of mass of the neural response toward the ST (Ensemble 1, High: fixation [4.25, –0.47] dva, perisaccadic [4.94, –0.61] dva; Low: fixation [5.14, –0.46] dva, perisaccadic [5.01, –0.84] dva. Ensemble 2, High: fixation [3.61, –0.97] dva, perisaccadic [4.15, –1.47] dva; Low: fixation [3.35, –1.08] dva, perisaccadic [4.09, –1.19] dva). We calculated the center of mass of neuronal responses relative to the ST across the nine probe locations, for High and Low trials for each neuron. A greater value means the center of mass is farther from the ST, while a smaller value means it is closer to the ST. For both monkeys, the center of mass of neuronal response over High trials shifts closer to the ST horizontally for probes presented perisaccadically compared with during fixation (Monkey 1: fixation = 4.74±0.12 dva, perisaccadic = 4.54±0.11 dva, *p* = 0.01; Monkey 2: fixation = 6.15±0.07 dva, perisaccadic = 5.98±0.05 dva, *p* = 0.01), whereas the center of mass of neuronal response over Low trials relative to the ST horizontally is not different for probes presented during the fixation and perisaccadic windows (Monkey 1: fixation = 4.48±0.12 dva, perisaccadic = 4.45±0.11 dva, *p* = 0.43; Monkey 2: fixation = 6.07±0.06 dva, perisaccadic = 5.98±0.05 dva, *p* = 0.12) (Fig. 3b). We then defined ‘center of mass shift’ as the perisaccadic center of mass subtracted from the center of mass during fixation, so a positive shift means shifting farther from the ST, and a negative shift means shifting closer to the ST. For both monkeys, there is no difference between the center of mass shift of High versus Low trials either horizontally (Monkey 1: High = –0.22±0.08 dva, Low = –0.07±0.07 dva, *p* = 0.25; Monkey 2: High = –0.20±0.07 dva, Low = – 0.13±0.07 dva, *p* = 0.40) or vertically (Monkey 1: High = –0.08±0.08 dva, Low = – 0.02±0.07 dva, *p* = 0.53; Monkey 2: High = –0.42±0.07 dva, Low = –0.52±0.06 dva, *p* = 0.59) (Fig. 3c).

**Figure 3.**
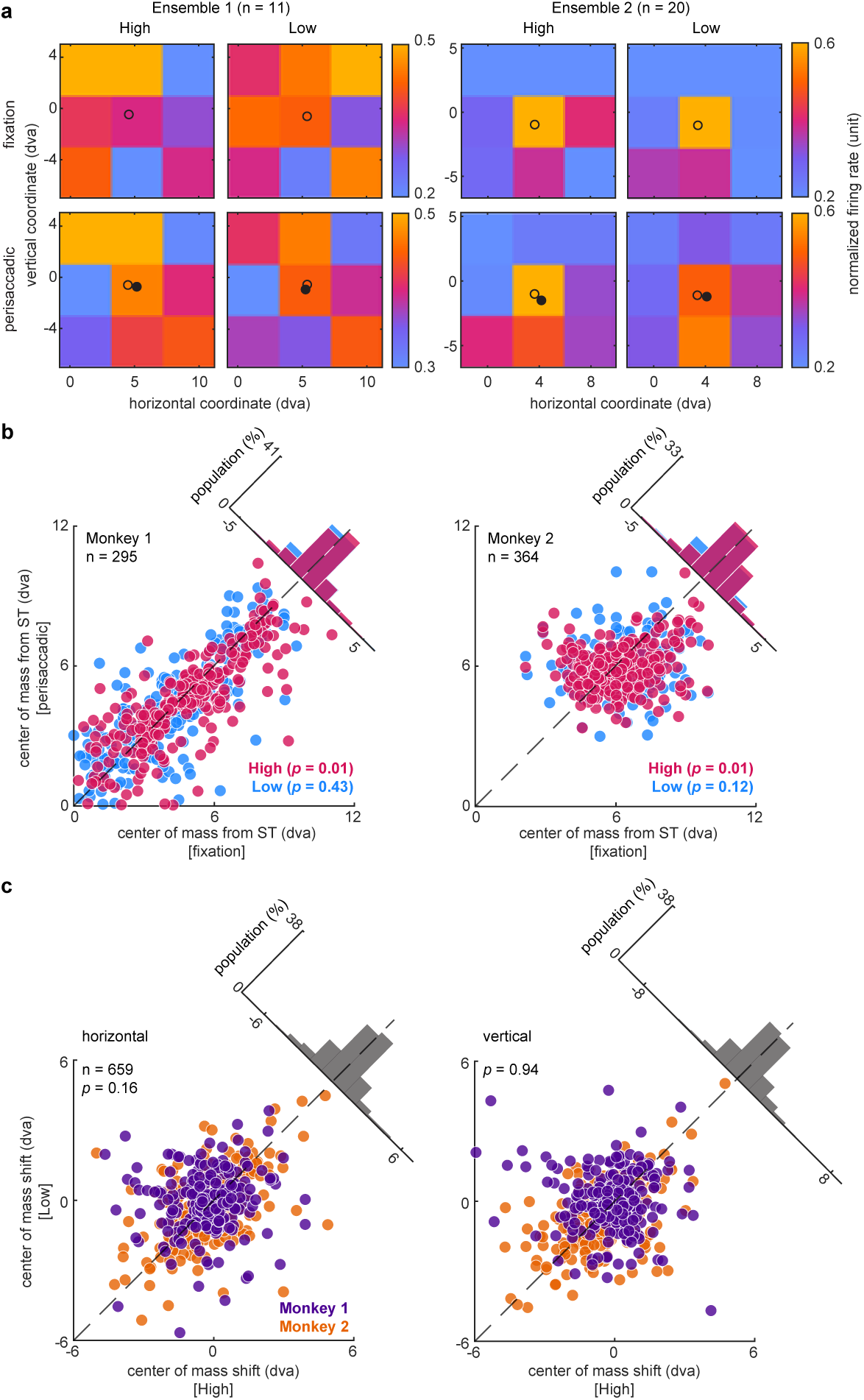
Neurons’ spatial sensitivity shifts toward the ST in trials with higher mislocalization magnitudes. **a.** Neuronal responses of two example ensembles of neurons (left and right sets of plots), for High and Low trials (left and right columns) at nine probe locations for probes presented during the fixation (top rows) and perisaccadic (bottom rows) windows. Both ensembles of neurons were recorded during sessions with rightward saccades toward the ST at [10, 0] dva. Color shows mean of normalized firing rates across neurons at each probe location. Circles show center of mass of normalized firing rates at nine probes during fixation (open circle) and perisaccadic (filled circle) windows. **b.** The center of mass of neuronal responses of High (magenta) versus Low (blue) trials. Plots show the center of mass of neuronal responses relative to ST horizontally for probes appearing during fixation (x-axis) versus perisaccadically (y-axis) for Monkey 1 (left; High: p = 0.01, Low: p = 0.43) and Monkey 2 (right; High: p = 0.01, Low: p = 0.12). **c.** The center of mass shift of neuronal responses relative to ST horizontally (left; p = 0.16) and vertically (right; p = 0.94) for High (x-axis) versus Low (y-axis) trials. ‘Center of mass shift’ is the perisaccadic center of mass subtracted from the center of mass during fixation. Positive shift means shifting farther from ST, and negative shift means shifting closer to ST. Plots show the combined population (n = 659 neurons) of Monkey 1 (purple) and Monkey 2 (orange). p-values were computed by two-sided Wilcoxon signed-rank tests. Histograms in the upper right show the distribution of differences.

To enhance the power of our analysis, we next focused on sessions and probes displaying the greatest and most consistent behavioral mislocalization. These were sessions with rightward saccades and visual probes located horizontally between 3 and 9 dva and vertically between –10 and 5 dva. We then repeated the analyses shown in Figure 1b–d and Figure 2 using only trials at the selected probe locations (Fig. 4). (We did not repeat the analysis in Figure 3 for the selected probes because that analysis requires all nine probes from each session to compute the center of mass.) Similar to Figure 1c-d, these probes show perisaccadic mislocalization (Fig. 4a) and localization error (Fig. 4b; Monkey 1: fixation = 0.45±0.19 dva, perisaccadic = −0.15±0.21 dva, *p* = 4.89e−09, Monkey 2: fixation = −0.61±0.13 dva, perisaccadic = −0.91±0.12 dva, *p* = 8.52e−05) opposite to the direction of the saccade. The modulation index of neurons at selected probes again demonstrates no difference between the High and the Low trials (Fig. 4c-d; Monkey 1: n = 146, High = 0.05±0.02, Low = 0.01±0.02, *p* = 0.24; Monkey 2: n = 456, High = 0.08±0.01, Low = 0.07±0.01, *p* = 0.43).

**Figure 4.**
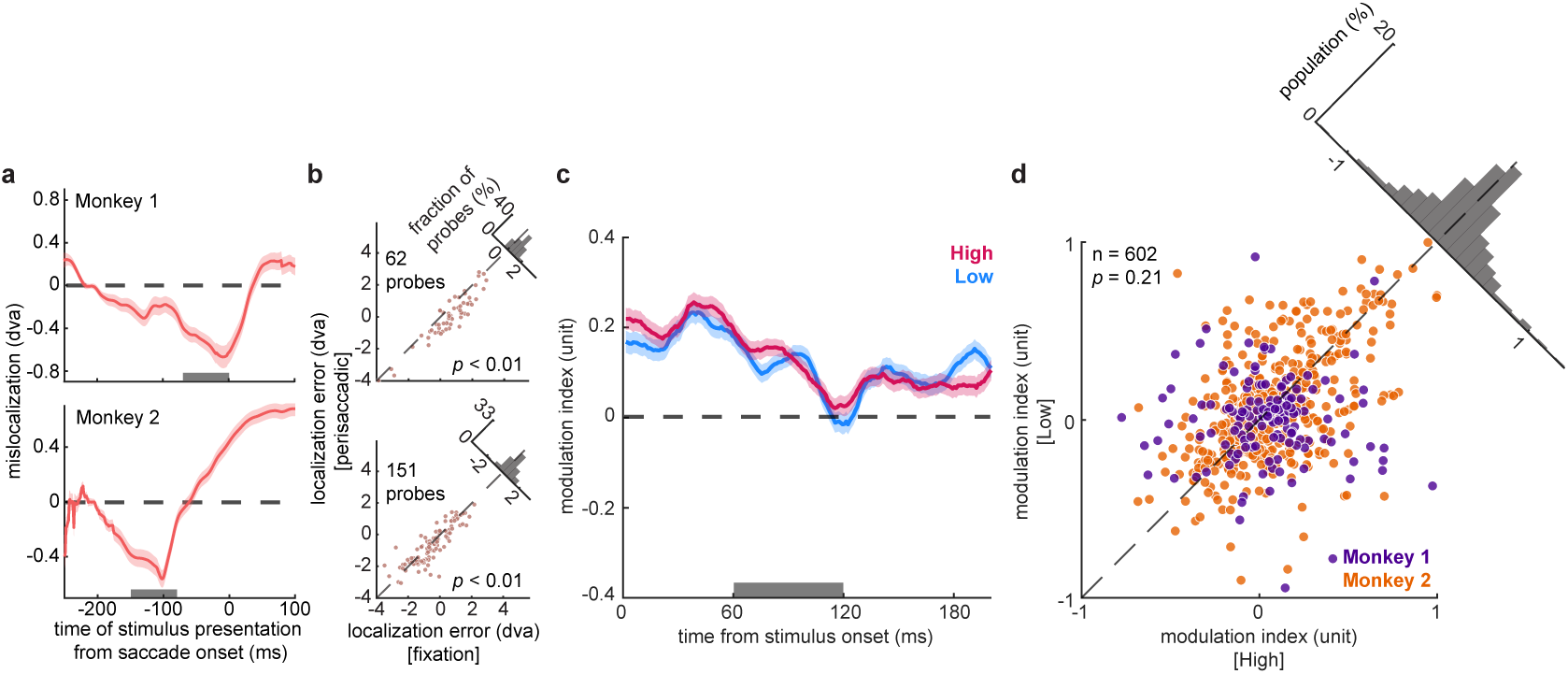
Behavioral effects and neuronal modulation for visual probes associated with most consistent behavioral mislocalization. **a.** Visual probes with horizontal locations within 3 to 9 dva and vertical locations within -10 to 5 dva were selected for the analyses shown in (a-d). The plot shows mean behavioral mislocalization (for probes in ±25 ms window), and the shaded area represents the SEM for Monkey 1 (top) and Monkey 2 (bottom). The number of probes included varies for different time points. Positive values indicate mislocalization in the saccade direction, and negative values reflect mislocalization opposite to the direction of the first saccade. The gray bar shows the perisaccadic time windows used for (b). **b.** Horizontal localization error for selected probes presented in fixation versus perisaccadic windows. Plots show horizontal localization error for probes appearing perisaccadically (y-axis) versus during fixation (x-axis) for Monkey 1 (top; *p* = 4.89e−09) and Monkey 2 (bottom; *p* = 8.52e−05). *p*-values were computed by two-sided Wilcoxon signed-rank tests. Histograms in the upper right show the distribution of differences. **c.** The modulation index of High trials (magenta) and Low trials (blue) over time from stimulus onset for selected probes. Mean ± SEM across neurons. The gray bar (60:120 ms) shows the analysis window used for (d). **d.** The mean modulation index within the analysis window for High versus Low trials for selected probes. Plots show the combined population of Monkey 1 (purple; n = 146 neurons, *p* = 0.24) and Monkey 2 (orange; n = 456 neurons, *p* = 0.43). *p*-values were computed by two-sided Wilcoxon signed-rank tests. Histograms in the upper right show the distribution of differences.

## Discussion

Precisely how different visual sensory areas contribute to the perception of stimulus location remains unclear. Around the time of saccades, the perceived location of stimuli changes (perisaccadic mislocalization); however, the link between perisaccadic mislocalization and saccade-related changes in neural responses has not been fully established. Previous studies have proposed that perisaccadic neural remapping may underlie mislocalization. Building on these theoretical frameworks, we designed and implemented a combined behavioral and electrophysiological framework to examine whether V4 neurons predict changes in spatial perception during saccades. We measured monkeys’ perception of stimulus location behaviorally and found perisaccadic mislocalization opposite to the saccade direction. We recorded from V4 neurons during the behavioral task and divided the trials into High and Low trials based on mislocalization magnitudes. We found no difference in neuronal modulations between the two groups. We found that the center of mass of neuronal responses for High trials shifts horizontally closer to the ST perisaccadically, but the shift is not statistically different from the Low trials. This lack of a relationship between V4 firing rate modulations and behavioral mislocalization across trials suggests that perceptual mislocalization may be governed by more complex neural coding mechanisms, beyond receptive field shifts based on the perisaccadic firing rate of V4 neurons.

We observed mislocalization opposite to the saccade direction for probes located between the FP and the ST, which is more consistent with a pattern of divergent than convergent mislocalization (Fig. 1a). However, because our paradigm did not include probes presented beyond the ST, we cannot determine whether the mislocalization pattern reflects a divergence or a uniform backward shift. Similarly, although we observed a sensitivity shift toward the ST, we cannot determine whether it reflects a uniform forward shift, as in future field remapping, or a convergent shift, as in ST remapping. Taken together, our findings are suggestive of the ‘unaware’ decoder hypothesis, but they do not establish a direct link between the behavioral mislocalization and the spatial shift in V4 neural sensitivity. Therefore, we conclude that the neural sensitivity shifts as measured by firing rate responses of V4 neurons is not sufficient to explain the observed perisaccadic mislocalization.

Studies have shown that future field remapping depends on corollary discharge^23,49^—an internal copy of the saccadic signal sent to other brain regions to inform them of the impending movement^50,51^. Corollary discharge provides the vector of the saccade that is available in advance of the upcoming saccade. Without access to the saccade vector prior to movement, it would not be possible to align the future RF with the target in the same way the current RF is aligned with the fixation point^52^. Recent theoretical studies propose that future field remapping based on corollary discharge and the ‘unaware’ assumption can explain the classic biphasic perisaccadic mislocalization^46,47,53^, where probes presented right before and around saccade onset are mislocalized in the saccade direction, and probes presented right after saccade onset are mislocalized opposite to the saccade direction^4,5,11^. However, whether corollary discharge is involved in ST remapping remains unclear. Studies have suggested that ST remapping is likely attention directed to the saccade target instead of the result of corollary discharge^46,47,52^. V4 neurons likely receive corollary discharge or attentional signal from upstream areas in the prefrontal cortex before the onset of a saccade, which results in a shift of RFs. Since our dataset was not explicitly designed to distinguish between these two forms of remapping, we cannot determine whether the observed sensitivity shift in neurons reflects future field remapping, ST remapping, or a combination of both (as suggested by previous reports^34–37^).

Our previous work showed perisaccadic spatial bias in extrastriate neurons around the ST and revealed the potential underlying neuronal response components^48^. In this study, we directly examined the link between behavioral mislocalization and neuronal responses recorded at the same time. Previously, we identified neuronal components relevant for spatial bias around −20:10 ms from saccade onset, and around 70:130 ms from stimulus onset, in neurons with RFs close to the ST. Most of the neurons analyzed in this study have RFs close to the ST, and we examined changes in neuronal sensitivity around 60:120 ms from stimulus onset in the perisaccadic window, similar to the computationally identified window. In both studies, we observed that neuronal sensitivity shifts horizontally toward the ST. In the present study, however, we found that the perisaccadic sensitivity shift of V4 firing rates was not related to the magnitude of behavioral mislocalization across trials. It is important to note that we predicted spatial bias in our prior study using kernels that represent the spatiotemporal properties of V4 neurons that trigger spikes, which are different from the stimulus-aligned firing rates used in this study. While we did not observe a trial-by-trial correlation between V4 perisaccadic firing rates and mislocalization magnitude, V4 may still contribute to mislocalization through more complex perisaccadic sensory representations, such as population dynamics, the temporal structure of responses, or interactions with other brain areas.

The frontal eye field (FEF), located in the prefrontal cortex, directly projects to V4 and is associated with both working memory and eye movements^54^. Literature has demonstrated that the FEF can modulate extrastriate responses related to behavior^55–57^ and that the FEF transmits working memory signals to visual areas, generating oscillations within V4 and influencing the timing of V4 spikes relative to these oscillation^58–61^. Future work can incorporate simultaneously recorded FEF signals to explore further how sensory and cognitive factors are integrated through inter-areal communication between the prefrontal and extrastriate cortices. This approach could deepen our understanding of the roles of attention and corollary discharge in driving the observed shifts in extrastriate sensitivity. It is also worth noting that the characteristics of the probe stimuli used in this study (brief, small, high-contrast) mean they are likely processed primarily through the magnocellular pathway. It is possible that perceived location depends more heavily on different neural regions based on the stimulus’s properties and task-relevant features (and so V4 responses might be more related to perceived location for colorful complex objects than for the probes used here). Simultaneous behavioral measurements and electrophysiological recordings from other areas (such as MT, MST, parietal, and prefrontal areas) will be needed to determine whether firing rate in any of these areas is more closely related to perceived location on individual trials. Together, these insights will bring us closer to understanding the neural basis of various facets of perception, and how they change around the time of eye movements.

## Methods

### Animals

All experimental procedures complied with the National Institutes of Health Guide for the Care and Use of Laboratory Animals and the Society for Neuroscience Guidelines and Policies. The protocols for all experimental, surgical, and behavioral procedures were approved by the Institutional Animal Care and Use Committees of the University of Utah.

We trained and recorded from two adult male rhesus macaques (*Macaca mulatta*; both 11 years old).

### Behavioral Paradigm

The behavioral task used in this study is called a Perisaccadic Localization Task. To start a trial, the monkey held fixation on a central FP. After the monkey had held fixation for 500 ms, an ST appeared 10 dva away from the FP horizontally. Each recording session had only one saccade direction (leftward or rightward). At a random time during fixation or saccade execution (1 to 301 ms after FP turns off), a 50-ms visual probe stimulus was presented in one of nine possible locations in a 3×3 grid. The distance between two adjacent probe locations is four dva in both the horizontal and vertical directions. After the monkey held fixation for 1000 ms, the FP disappeared, which was the go cue to saccade to the ST. The monkey then made a second saccade to report his perceived location of the probe stimulus. If the monkey reported within 4 dva radius of the correct location, the stimulus reappeared as feedback, and the monkey received a juice reward. The FP and ST were white circles (full contrast) with a radius of 0.25 dva, and the probe stimuli were red circles with a radius of 0.4 dva against a black background. We collected behavioral data during 91 sessions for Monkey 1 and 81 sessions for Monkey 2.

### Electrophysiology

The Perisaccadic Localization Task is based on the tasks used in previous behavioral studies to characterize perisaccadic mislocalization in monkeys^6,8,14^; it was designed to be similar to paradigms we used in the past for other neurophysiological studies in terms of ambient lighting conditions, probe size, background color, and saccade amplitude. During each neurophysiological recording session, the grid of possible probe locations was placed around the estimated presaccadic RF centers of the neurons recorded (estimated RF position based on audible responses to sweeping bar stimuli). We used single tungsten microelectrodes of 200 μm diameter, with epoxylite insulation (FHC, Bowdoin, ME) and linear 16-channel array electrodes (U-Probe and S-Probe, Plexon Inc., Dallas, TX; Cheetah v5.7.4 in Neuralynx acquisition systems) at a sampling rate of 32 kHz. Electrodes were inserted using a hydraulic microdrive (Narishige, Japan). Neural waveforms were sorted offline using the Plexon offline spike sorter. A total of 494 neurons from Monkey 1 and 509 neurons from Monkey 2 were used in this study.

### Eye-tracking

While the monkey was performing the task, we monitored its eye movements with an infrared optical eye-tracking system (EyeLink 1000 Plus Eye Tracker, SR Research Ltd., Ottawa, CA) with a resolution of <0.01 dva (based on the manufacturer’s technical specifications), and a sampling frequency of 2 kHz. The presentation of the visual stimuli on the screen was controlled using the MonkeyLogic toolbox.

### Trial-by-trial Analysis

Throughout the paper, mislocalization is defined as the horizontal distance between the reported location of each stimulus when presented around the time of the first saccade (Monkey 1: −70:0 ms; Monkey 2: −150:−80 ms), compared with the average reported location for stimuli presented during fixation (−250:−180 ms). For each trial, mislocalization was calculated by subtracting the mean reported location across all fixation trials from the reported location of that trial. Localization error was defined as the distance between the reported location of each stimulus and the actual location of the probe presented for each trial. We used two-sided Wilcoxon signed-rank tests to report *p*-values for all our statistical comparison analyses.

High trials were defined as the trials associated with mislocalization magnitudes (absolute values of mislocalization) in the top 45^th^ percentile for each probe location in a session, and Low trials are the ones associated with mislocalization magnitudes in the bottom 45^th^ percentile. The modulation index in Figure 2 was computed using trials from 88 sessions of Monkey 1 (High: fixation = 1.33±0.04 dva, perisaccadic = 1.79±0.04 dva; Low: fixation = 0.31±0.01 dva, perisaccadic = 0.53±0.02 dva; mean mislocalization magnitude ± SEM) and 60 sessions of Monkey 2 (High: fixation = 0.41±0.19 dva, perisaccadic = 1.73±0.78 dva; Low: fixation = 0.19±0.02 dva, perisaccadic = 0.78±0.07 dva). The modulation index in Figure 4c-d was computed using trials from 23 sessions of Monkey 1 (High: fixation = 1.36±0.07 dva, perisaccadic = 1.58±0.07 dva; Low: fixation = 0.31±0.02 dva, perisaccadic = 0.44±0.02 dva; mean mislocalization magnitude ± SEM) and 58 sessions of Monkey 2 (High: fixation = 0.34±0.04 dva, perisaccadic = 0.89±0.03 dva; Low: fixation = 0.14±0.01 dva, perisaccadic = 0.42±0.02 dva).

### Center of Mass Analysis

The center of mass (*d_C_*) of neuronal responses (Fig. 3; Supplementary Fig.1) was calculated using Equation (2):

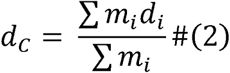

where *m_i_* represents the mean neuronal firing rates 60:120 ms from stimulus onset at probe *i*, and *d_i_* represents the coordinates of probe *i* on screen in sample neurons (panel a), or the horizontal coordinate of probe *i* relative to the ST location in the population of neurons (panel b).

## Data Availability

The datasets generated and/or analyzed for this study will be available on a public repository upon the acceptance of the manuscript.

## Acknowledgements

The authors would like to thank the animal care personnel at the University of Utah. We specifically thank Rochelle D. Moore and Dr. Tyler Davis for their assistance with the NHP experiments. This work was supported by NIH EY026924 and NIH NS113073 to B.N.; NIH EY031477 and AFOSR FA9550-24-1-0235 to N.N.; NIH EY014800 and an Unrestricted Grant from Research to Prevent Blindness, New York, NY, to the Department of Ophthalmology & Visual Sciences, University of Utah.

## Author Contributions

N.N. and B.N. conceived the study. B.N. performed the surgical procedures. N.N., B.N., and G.W. designed the experiment. N.N., B.N., G.W., and K.C. designed the analysis. G.W. performed the physiology experiments and acquired data. G.W. performed the data analysis. G.W., K.C., and N.N. wrote the manuscript.

## Competing Interests

The authors declare no competing interests.

**Supplementary Information** is available for this paper.

