## Supplementary Information for "Limits of V4 perisaccadic firing rate modulations in explaining perceptual mislocalization"

Behrad Noudoost

Neda Nategh

**Word counts**

324

**Number of figures**

2

Supplementary Figures

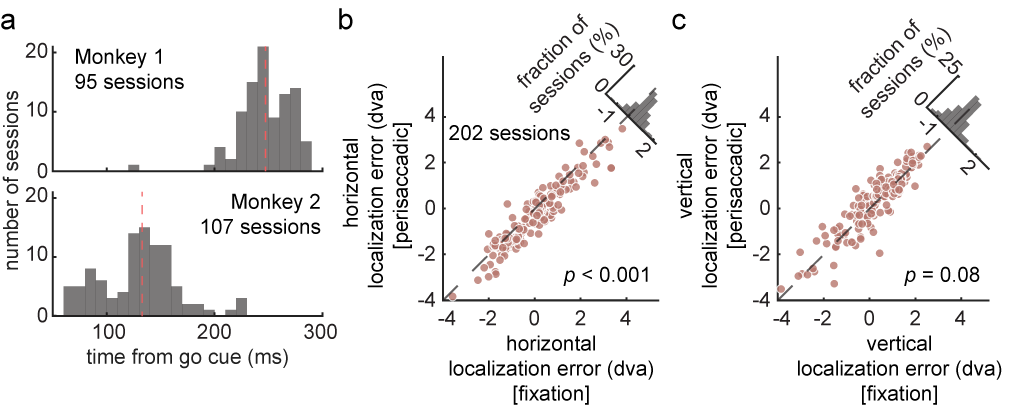

**Supplementary Figure 1. Reaction time difference of two monkeys, and localization errors in horizontal vs. vertical directions. a.** Histograms of reaction time of two monkeys. Reaction time was measured as the time of the first saccade onset from go cue (fixation point offset; see Fig. 1b). Red dashed lines show the median reaction time (Monkey 1: median = 247.21 ms, Monkey 2: median = 132.87 ms). **b**. Horizontal localization error for each of 202 sessions, combining sessions for both Monkey 1 and Monkey 2. Plot shows mean horizontal localization error of each session for probes presented in the fixation vs. perisaccadic windows (*p* = 8.21e-17). **c.** Vertical localization error of each of 202 sessions for probes presented in the fixation vs. perisaccadic windows (*p* = 0.08). We only measured the vertical localization errors for probes below the ST to avoid averaging out potential compression effects. Histograms in the upper right of all panels show the distribution of differences.

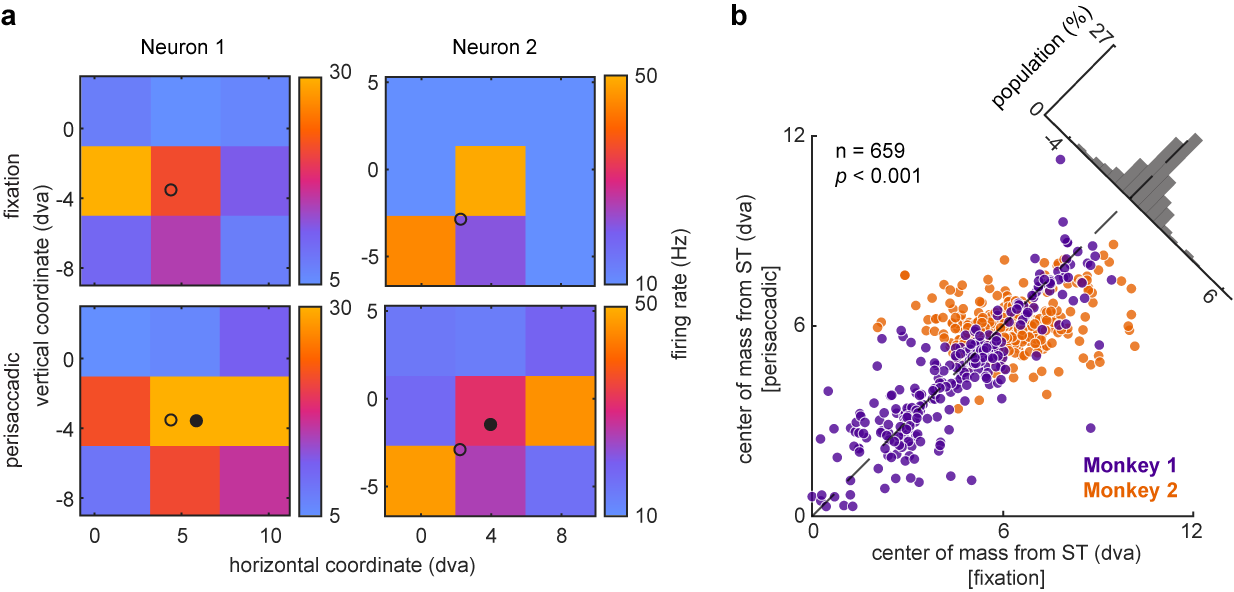

**Supplementary Figure 2. Spatial sensitivity shifts in sample neurons and population. a.** Neuronal responses of two sample neurons (left and right) at nine probe locations for probes presented during the fixation (top) and perisaccadic (bottom) windows. Both neurons were recorded during sessions with rightward saccades toward the ST at [10, 0] dva. Plot shows firing rates of the neurons and center of mass of firing rates at nine probes during fixation (open circle; Neuron 1: [4.39, −3.52] dva, Neuron 2: [2.22, −2.84] dva) and perisaccadic (filled circle; Neuron 1: [5.83, −3.63] dva, Neuron 2: [3.97, −1.41] dva) windows. **b.** The center of mass of neuronal responses during fixation vs. perisaccadic time windows. Plots show the center of mass of neuronal responses relative to ST horizontally for probes appearing during fixation (x-axis) vs. perisaccadically (y-axis) (n = 659 neurons, p = 2.99e-4). Plot shows the combined population of Monkey 1 (purple; n = 295, p = 0.02) and Monkey 2 (orange; n = 364, p = 2.14e-3).
